# Quantification and statistical modeling of Chromium-based single-nucleus RNA-sequencing data

**DOI:** 10.1101/2022.05.20.492835

**Authors:** Albert Kuo, Kasper D. Hansen, Stephanie C. Hicks

**Affiliations:** Department of Biostatistics, Johns Hopkins Bloomberg School of Public Health, Baltimore, Maryland, USA; Department of Genetic Medicine, Johns Hopkins School of Medicine, Baltimore, Maryland, USA

**Keywords:** gene expression, single-cell RNA-sequencing, single-nucleus RNA-sequencing, zero-inflation, negative binomial distribution, Poisson distribution, binomial distribution

## Abstract

In complex tissues containing cells that are difficult to dissociate, single-nucleus RNA-sequencing (snRNA-seq) has become the preferred experimental technology over single-cell RNA-sequencing (scRNA-seq) to measure gene expression. To accurately model these data in downstream analyses, previous work has shown that droplet-based scRNA-seq data are not zero-inflated, but whether droplet-based snRNA-seq data follow the same probability distributions has not been systematically evaluated. Using pseudo-negative control data from nuclei in mouse cortex sequenced with the 10x Genomics Chromium system, we found that snRNA-seq data follow a negative binomial distribution, suggesting that parametric statistical models applied to scRNA-seq are transferable to snRNA-seq. Furthermore, we found that the quantification choices in adapting quantification mapping strategies from scRNA-seq to snRNA-seq can play a significant role in downstream analyses and biological interpretation. In particular, reference transcriptomes that do not include intronic regions result in significantly smaller library sizes and incongruous cell type classifications. We also confirmed the presence of a gene length bias in snRNA-seq data, which we show is present in both exonic and intronic reads, and investigate potential causes for the bias.

## INTRODUCTION

Single-nucleus RNA-sequencing (snRNA-seq) is a common experimental technology to profile gene expression in frozen cells or cells that are hard to dissociate, such as in brain tissue (Lake et al., 2016; Slyper et al., 2020). Previous studies have shown that snRNAseq offers substantial advantages over single-cell RNA-sequencing (scRNA-seq), including reduced dissociation bias (Habib, Li, et al., 2016; Bakken et al., 2018) and the ability to capture rare cell types (Wu et al., 2019). However, several questions remain on the degree to which existing tools used to analyze scRNA-seq data can be used in application of snRNA-seq data, including, (i) what is an appropriate genomic unit (e.g. exonic regions, intronic regions, etc) to quantify reads for down-stream analysis and (ii) what are appropriate probability distribution(s) to model measurement error or noise. We begin by discussing these two topics in greater detail.

A standard approach to remove ribosomal RNA (rRNA) from scRNA-seq and snRNA-seq protocols, such as droplet-based technologies with unique molecular identifiers (UMIs) from the 10x Genomics Chromium system (Zheng et al., 2017), is to select polyadenylated RNA (polyA) transcripts using oligo (dT) primers. During the process of transcription, the gene is converted into a precursor mRNA (referred to as pre-mRNA) that contains both exonic and intronic regions. Mature mRNA is formed after the intronic regions have been spliced out of pre-mRNA, leaving only exonic regions. RNA processing happens in the nucleus so we expect mRNA existing outside the nucleus to be without introns, which have been experimentally verified (Cooper, Hausman, 2007; Ding et al., 2020; Lee et al., 2020). Hence, raw sequencing reads from scRNA-seq protocols are typically quantified using reference genomes or transcriptomes with only exonic regions, which has been shown to be sufficient for downstream analyses such as accurately classifying cell types (Habib, Avraham-Davidi, et al., 2017; Bakken et al., 2018; Ding et al., 2020). However, it is unclear whether it is also sufficient to quantify reads from snRNA-seq protocols to only exonic regions for downstream analyses. If not, then how should intronic regions be incorporated, that is, whether reads should be quantified with exonic and intronic regions separately or mapped to full-length spliced and unspliced transcripts (pre-mRNA) in snRNA-seq data. For example, previous work has shown these choices need to be carefully considered for RNA velocity with scRNA-seq (Soneson et al., 2021) and may be also necessary for obtaining high-quality results downstream with snRNA-seq data (Bakken et al., 2018).

Furthermore, when considering quantified reads from intronic regions with snRNA-seq data, it has been previously suggested that there is a gene length bias (Chamberlin, Quinlan, 2020). The source of this bias may lie with the enriched pre-mRNA in snRNA-seq data, as scRNA-seq data sequenced under UMI-based protocols, such as CEL-Seq, SMARTer, and CEL-Seq with InDrop, are not believed to exhibit a length bias (Phipson et al., 2017); we note this has not been specifically investigated with the 10x Genomics Chromium system. However, the extent to which reads from intronic versus exonic regions contribute to the length bias and the potential causes of the bias are not well understood. In addition, it is unclear whether the length bias is a function of the full length of the unspliced transcript, consisting of both exonic and intronic regions, or the length of the spliced transcripts, consisting of exonic regions.

Next, given that an appropriate unit of quantification has been determined, another question is the choice of probability distributions to model snRNA-seq data. Much progress has been made on investigating the appropriateness of distributions to model scRNA-seq data (Hafemeister, Satija, 2019; Townes et al., 2019; Svensson, 2020; Choi et al., 2020; Ahlmann-Eltze, Huber, 2021; Sarkar, Stephens, 2021; Jiang et al., 2022; Choudhary, Satija, 2022) and an open question is whether similar distributions can be used to model measurement error or noise in Chromium-based scRNA-seq data. In the context of scRNA-seq, previous work has argued that intronic reads represent experimental and transcriptional noise (Harati et al., 2014) and are not usable in gene quantification (Zhao et al., 2018). As snRNA-seq enriches for transcripts in the nucleus with both mRNA and pre-mRNA, and scRNA-seq enriches for mostly mature mRNA, it is unclear if the measurement error in observed in snRNA-seq data is likewise affected by the inclusion of the intronic reads.

Furthermore, the consequences on the choice of appropriate probability distributions to model measurement error are not well understood. Previous work has demonstrated that Chromium-based scRNA-seq data with UMIs are not zero-inflated and can be accurately modeled using Poisson, negative binomial, or multinomial distributions (Townes et al., 2019; Hafemeister, Satija, 2019; Svensson, 2020). This was demonstrated using negative control data, where no biological heterogeneity is expected, by adding a controlled amount of RNA to each droplet. However, to the best of our knowledge, there has been no comparable analysis for the analysis of Chromium-based snRNA-seq data while mapping reads to both introns and exons.

In this paper, we first evaluate the choice of probability distributions to model measurement error in snRNA-seq data by creating pseudo-negative control datasets. Next, we evaluate reference transcriptomes, which differ in how to include exonic and intronic regions, used in quantification mapping tools and consider the impact on cell type classification in downstream analyses. Then, we investigate and confirm the existence of a gene length bias in both intronic and exonic reads.

## RESULTS

Throughout, we used snRNA-seq data from two mouse cortices (Ding et al., 2020) measured on the 10x Genomics Chromium platform (Zheng et al., 2017). We begin by creating pseudo-negative control datasets (**Figure 1**) by working with subsets of mouse cortex cell types to identify more homogenous populations of cells, where we expect less biological variation within a cell type than across cell types (**Figure S1**). Throughout, we use *italicized* font when referring to datasets or different types of reference transcriptomes, referred to as *transcripts, preandmrna, introncollapse*, and *intronseparate*. These reference transcriptomes differ in how to include exonic and intronic regions used in the `salmon alevin` (Srivastava et al., 2019) tool to perform quantification mapping of raw sequencing reads.

**Figure 1.**
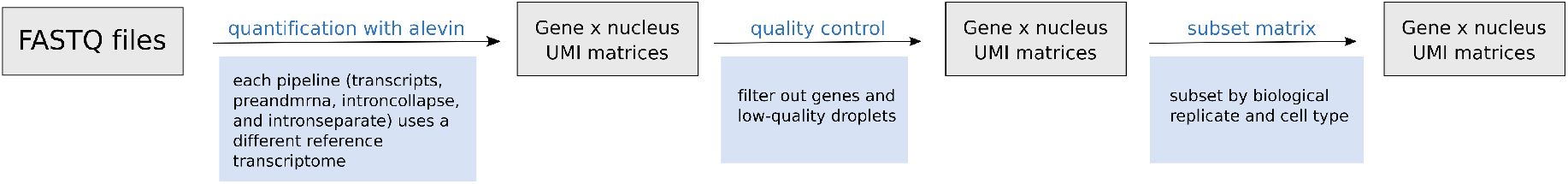
Overview of computational workflow to create pseudo-negative control snRNA-seq data. Raw sequencing reads (FASTQ files) from *N* = 2 mouse mouse cortices (Ding et al., 2020) were processed with `salmon alevin` to perform quantification mapping with four ways for how to include exonic and intronic regions in a reference transcriptome, referred to as (i) *transcripts*, (ii) *preandmrna*, (iii) *introncollapse*, or (iv) *intronseparate* (see **Table S1** for details). This results in four matrices of unique molecular identifier (UMI) counts with genes along the rows and nuclei along the columns. Next, we apply quality control metrics to filter out low-quality nuclei and lowly expressed genes. Finally, we stratify the nuclei by cell type (*C* = 7) and biological replicate (*N* = 2). Pseudo-negative control data represents analyses performed on these stratified subsets where we expect less biological variation within a cell type than across cell types.

### Chromium-based single-nucleus RNA-seq data are not zero-inflated

We begin by considering a pseudo-negative control dataset made with the *preandmrna* reference transcriptome, which uses mRNA and pre-mRNA (**Table S1**). However, at the end of this section, we show consistent results with the other reference transcriptomes. Here, we use the pseudo-negative control data to investigate whether the same probability distributions used to model measurement error in Chromium-based scRNA-seq can be used for Chromium-based snRNA-seq data.

Common distributions to model measurement error in scRNA-seq with UMI counts include the binomial, Poisson, and negative binomial (NB) distributions (Grün et al., 2014; Vieth et al., 2017; Svensson, 2020; Choudhary, Satija, 2022). Historically, it has been argued the scRNA-seq are “zero-inflated”, where the fraction of observed zeros in the single-cell counts is larger than what is expected under a specific distribution, such as the NB (Pierson, Yau, 2015; Risso et al., 2018; Jiang et al., 2022). The NB distribution has two parameters: a mean (or rate) parameter (*µ*) and a dispersion parameter (*ϕ*). This dispersion parameter can be estimated as an overall dispersion parameter for a given dataset or it can be estimated for each gene (or feature). Historically in application of bulk RNA-sequencing data, the interpretation of the *ϕ* parameter is to quantify how much extra biological variation is observed on top of technical variation (Robinson et al., 2010). However, in our application of snRNA-seq negative control data, we aim to estimate the parameters *µ* and *ϕ* to investigate if the NB distribution can be used to accurately capture the measurement error or technical variation observed from snRNA-seq data. For a random variable *X* that follows a NB with a given *µ* and a dataset-specific *ϕ*, the variance is

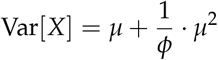

and the probability of observing a zero count is given by

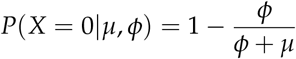

For a given cell type, we can compute for each gene (1) the empirical mean and empirical variance (commonly referred to as the “mean-variance relationship”) (Chen et al., 2014) and (2) the empirical mean and observed fraction of zero counts. We compare these observed quantities, the empirical variance and the observed fraction of zero counts, to what we expect them to be under the binomial, Poisson, and NB distributions. We also consider a NB distribution with a gene-specific dispersion parameter *ϕ*. Considering all four of these distributions, we calculate the Bayesian information criterion (BIC) (Burnham, Anderson, 2004) to assess the best model fit, where we sum up the log-likelihood across genes and assume the genes are independent. Lastly, we create quantile-quantile plots from a Pearson’s goodness-of-fit test under the Poisson model to assess the fit of the Poisson distribution to the observed counts.

Using these pseudo-negative control datasets with the excitatory neurons (**Figure 2a-d**), inhibitory neurons (**Figure 2e-h**) and astrocytes (**Figure 2i-l**), we found the empirical behavior of Chromium-based snRNA-seq UMI counts is closely approximated by standard probability distributions (**Figure 2a,e,i**) and are not zero-inflated (**Figure 2b,f,j**). Most genes exhibit variances and fraction of zero counts that can be approximated by a binomial or Poisson distribution, but the NB distribution provides the best fit using BIC (**Figure 2c,g,k**). As noted in Figure 2a-b, the binomial and Poisson distributions provide nearly indistinguishable theoretical fits, but differ from the NB distribution at higher empirical means. However, the difference in BIC between the negative binomial distribution and the binomial or Poisson distribution is driven primarily by a few genes (**Supplementary Figure S10**). The largest BIC values, and therefore the poorest fit, was found for the NB distribution with gene-specific overdispersion parameters (“G-S negative binomial”), which indicates that the gain in likelihood from having an overdispersion parameter for every gene is outweighed by the BIC penalty on the increase in the number of parameters. Using a Pearson’s goodness-of-fit test under a Poisson model, we also found that the gene counts exhibit deviations from the Poisson distribution (**Figure 2d,h,i**), which further suggest that the Poisson distribution does not adequately capture the technical variation in the counts. In addition to cell types, we found that these results also hold true across different biological replicates (the two mouse cortices), and reference transcriptomes (**Figures S2, S3, S4; Table S2**).

**Figure 2.**
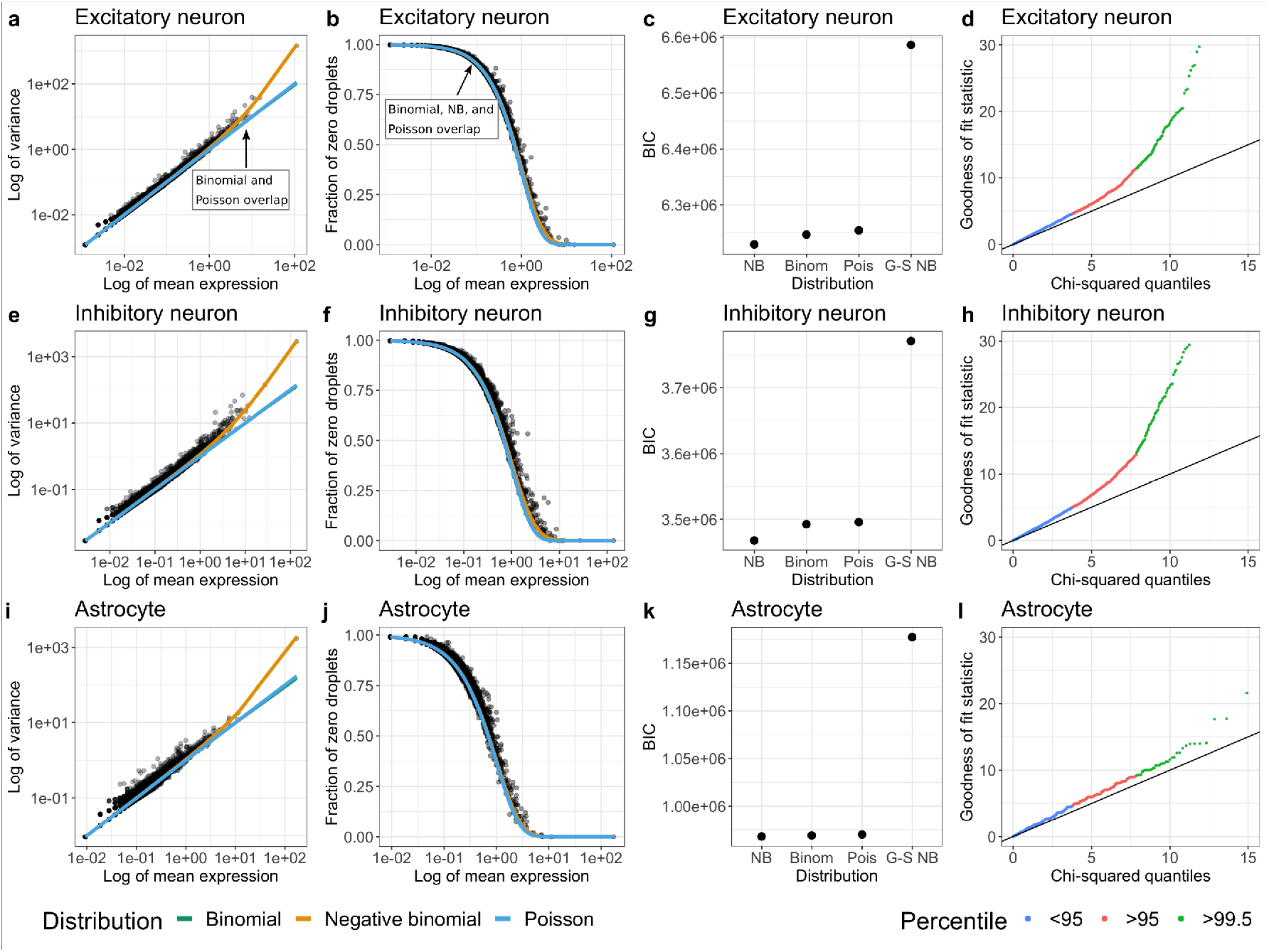
Chromium-based single-nucleus RNA-seq data is not zero-inflated. Using subsets of cell types, **(a-d)** Excitatory neurons, **(e-h)** Inhibitory neurons, and **(i-l)** Astrocytes, from Cortex 1 in the mouse cortex dataset (Ding et al., 2020), the leftmost column shows the log-transformed empirical mean (*x*-axis) and variance (*y*-axis) for each gene (black dots) with the theoretical variance (colored lines). The second column shows the log-transformed empirical mean (*x*-axis) and observed fraction of zeros (*y*-axis) for each gene (black dots) with the expected fraction of zeros under each distribution (colored lines). The third column shows BIC value across all genes (assuming the genes are independent) for each distribution: negative binomial (NB) with one overdispersion parameter estimated for the dataset, binomial (Binom), Poisson (Pois), and NB with gene-specific overdispersion parameters (G-S NB). The rightmost column shows the quantile-quantile plot under the Poisson model with the theoretical chi-squared quantile on the *x*-axis and the observed chi-squared statistic on the *y*-axis. Reads were quantified with the *preandmrna* reference transcriptome.

### Reference transcriptomes with intronic regions increases the total number of mapped reads

We continue with the same snRNA-seq data from the mouse cortex (Ding et al., 2020), but here we consider four reference transcriptomes (*transcripts, preandmrna, introncollapse*, and *intronseparate*), which differ in how to include exonic and intronic regions, used for quantification mapping with the `salmon alevin` (Srivastava et al., 2019) tool. The *transcripts* reference uses only the spliced transcripts as target sequences, while the other three quantification references additionally incorporate intronic regions as target sequences in different ways (**Table S1**). We evaluate how the choice of the reference transcriptome can impact the total number of mapped reads, and the number of mapped reads to subsets of genes including protein coding genes and pseudogenes. We found that the library size for the *transcripts* reference was smaller than the library sizes for the other three references (**Figure 3a**). We observe a similar disparity between references for the number of reads in protein-coding genes (**Figure 3b**). Interestingly, we found a decrease in the number of reads mapping to processed pseudogenes in the references with intronic reads (**Figure 3c**). This may occur if true pre-mRNA reads are mapped to other regions like processed pseudogenes in the *transcripts* reference due to the lack of intronic target sequences, but are correctly mapped to the intronic regions of genes in the other references. However, only minor differences in the number of mapped reads appear between the three references that incorporate intronic regions (*preandmrna, introncollapse*, and *intronseparate*). Finally, we also found an increase mapped reads to long non-coding RNA and antisense for the references that included intronic regions (**Figure S5**). This suggests that for snRNA-seq, the primary difference with respect to total mapped reads is driven by the incorporation of intronic regions in target sequences or not, rather than the specific ways in which they are incorporated.

**Figure 3.**
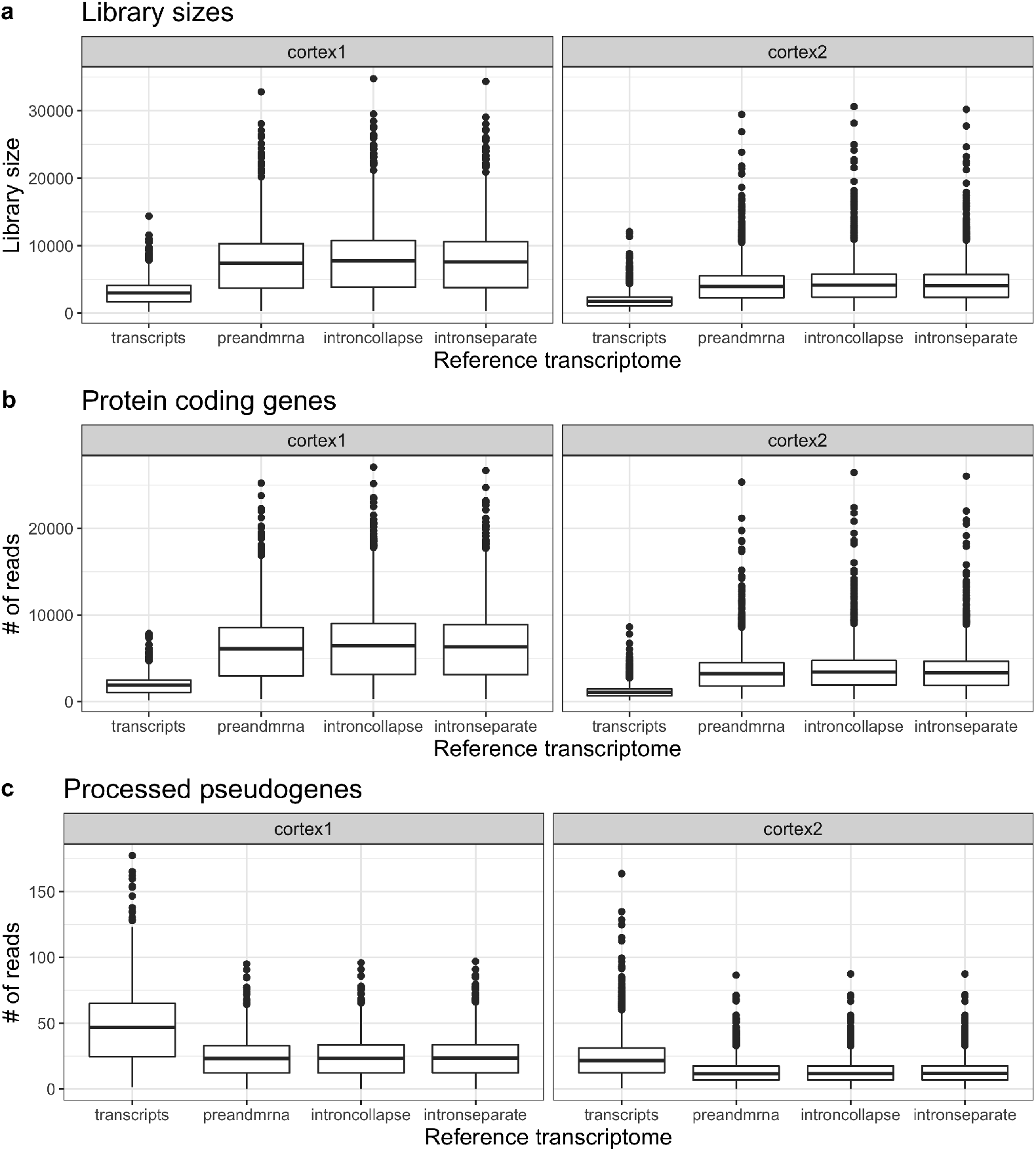
Incorporating intronic regions into reference transcriptomes leads to larger total mapped reads and reads mapping to protein coding genes. Left and right columns represent two biological replicates (mouse Cortex1 (left) and Cortex2 (right)) that were sequenced and quantified using four reference transcriptomes (does not include intronic reads: *transcripts*; includes intronic reads: *preandmrna, introncollapse, intronseparate*). Boxplots of the number of **(a)** total UMIs for each nuclei (or library sizes), **(b)** reads mapped to protein-coding genes, and **(c)** reads mapped to processed pseudogenes.

### Reference index impacts cell type classification

An example of how the lower mapping rate in the *transcripts* reference affects downstream analysis is in appli-cation of cell type classification. To demonstrate this, we used the reference-based cell type annotation algorithm `SingleR` (Aran et al., 2019) to identify cell types using each of the four reference transcriptomes. We compared these cell type classification labels to the labels provided by the authors of the original paper (Ding et al., 2020) (**Figure 4a**). We found that for the most common cell types (as classified by Ding et al., 2020), there is a high level of agreement with the `SingleR` cell type classification across most refence transcriptomes. However, we observe a higher discordance to the cell type classified by Ding et al., 2020 in the *transcripts* reference, especially for certain cell types. For instance, among the nuclei labeled as astrocytes by the authors, more nuclei are labeled as quiescent neural stem cells (qNSCs) instead of astrocytes by `SingleR` in the *transcripts* reference compared to the others (*preandmrna, introncollapse*, and *intronseparate*). Because `SingleR` is a reference-based algorithm, it depends on reads mapping to known marker genes to accurately classify cell types. Upon further inspection of the marker genes for astrocytes and qNSCs, we found more reads mapping to astrocyte marker genes compared to qNSC marker genes for only the references that include intronic reads, among the nuclei labeled as astrocytes by the authors (**Figure 4b**). In contrast, using the *transcripts* reference, the ratio of counts in astrocyte marker genes compared to qNSC marker genes is close to 1 on average for the astrocyte nuclei, resulting in some of these nuclei being classified as astrocytes and others classified as qNSCs by `SingleR`. This result indicates that the increase in mapping rate with the inclusion of intronic reads, as described in the previous section, does not occur uniformly across all marker genes, and real biological signal may be lost when intronic reads are not included.

**Figure 4.**
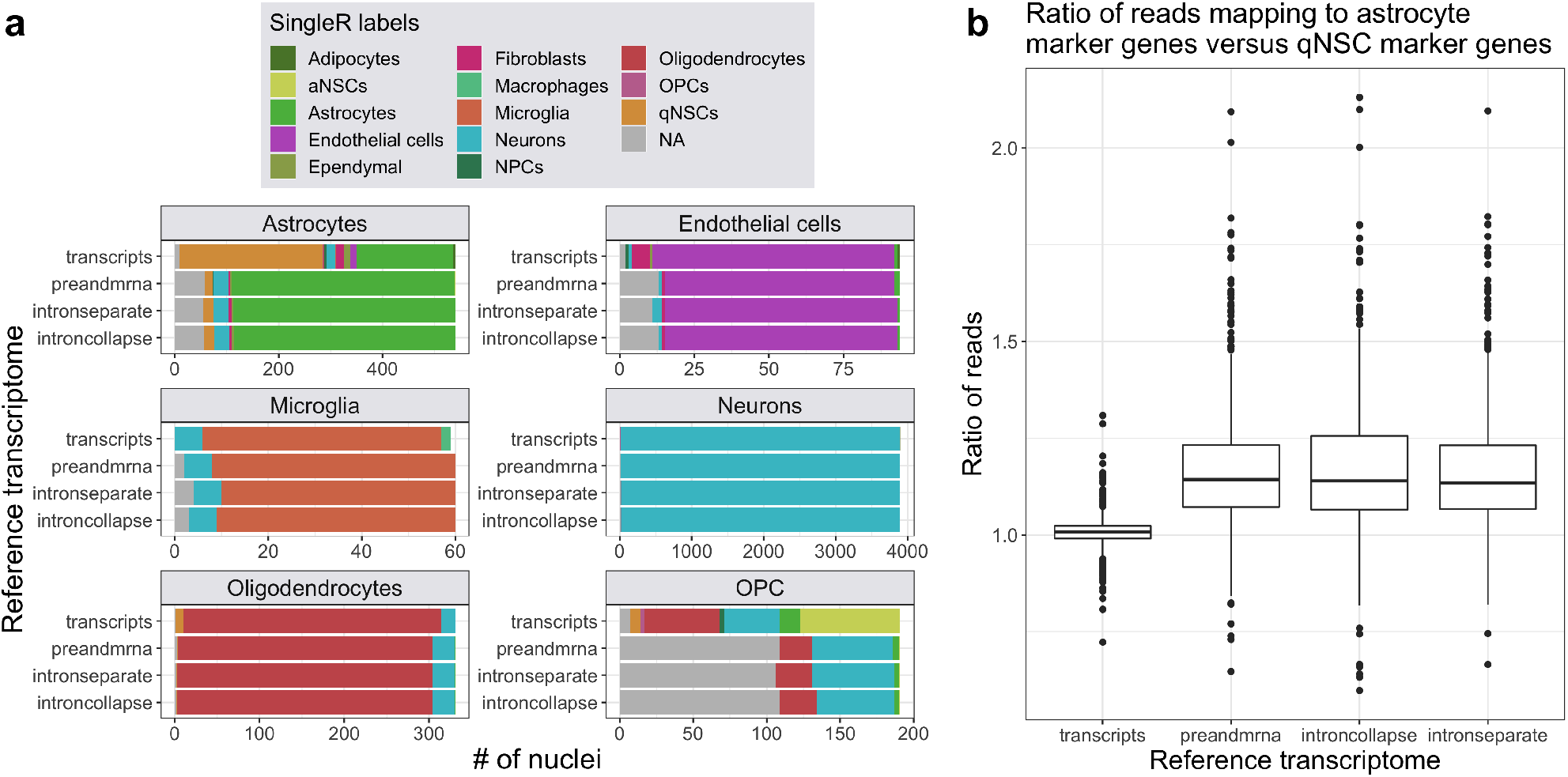
Choice of reference index can impact the classification of cell types in scRNA-seq data (a) For each cell type (labeled facets, classified by Ding et al., 2020), the bar plots show the number of nuclei that are assigned to different cell types by the reference-based `SingleR` annotation algorithm in each reference (*transcripts, preandmrna, introncollapse*, or *intronseparate*). Excitatory neurons and inhibitory neurons are combined into one cell type named ‘Neurons’ as the training dataset used in `SingleR` does not distinguish between them. **(b)** For the nuclei classified as astrocytes by Ding et al., 2020, the ratio of UMI counts in astrocyte marker genes to UMI counts in qNSC marker genes is elevated in reference indices that incorporate intronic regions (*preandmrna, introncollapse, intronseparate*) versus those that do not (*transcripts*).

We also observed major differences in cell type classification between references for the oligodendrocyte progenitor cell (OPC) cell type. In the *transcripts* reference, most of the OPC nuclei are classified into one of several cell types, while with the other references, nearly half of the OPC nuclei were not assigned a label. Cells are not assigned a label by `SingleR` when there is not enough signal to unambiguously assign a cell type classification, for example, when a given cell or nuclei has an expression profile equally similar to two or more cell types (Aran et al., 2019). Therefore, we observe that quantification choices can also influence cell type classification in less apparent ways, namely by determining whether a cell type label is assigned or not.

Similar to the number of mapped reads, the major differences in cell type classification remain between the choice of reference transcriptome to include intronic regions or not, with more minor differences among using references that include intronic regions. This shows that the specific ways of defining intronic regions in the reference transcriptome is less consequential than the choice to include or not include intronic regions.

### Chromium-based snRNA-seq data exhibit a gene length bias

In this section, we continue with the same snRNA-seq data (Ding et al., 2020), but here we show that there is a gene length bias, namely we observe a higher level of expression for genes that are longer compared to genes that are shorter. This is surprising because it is assumed that scRNA-seq and snRNA-seq UMI-based protocols, unlike full-length transcript protocols, do not exhibit a gene length bias due to the polyA selection on the 3’ end of the mRNA molecule (Phipson et al., 2017; Vallejos et al., 2017; Zheng et al., 2017). Nevertheless, a length bias has been previously described in snRNA-seq data (Chamberlin, Quinlan, 2020) and has been suggested to be caused by internal poly-A priming (below).

We begin by grouping genes into ten bins by their preandmrna length, with each bin containing the same number of genes. The ‘preandmrna’ length refers to the full gene length, which includes both the intronic and exonic regions of a gene (see **Supplementary Figure S6a** for a comparison between ‘transcript’ length and ‘preandmrna’). Most importantly, we observe a gene length bias, where the number of total counts per gene (sum of counts across all nuclei) increases with the gene length using both the *transcripts* and *preandmrna* references (**Figure 5**). The bias is stronger in the *preandmrna* reference, which suggests that the intronic reads play a major role in the length bias. We observe a similar trend using the *introncollapse* and *intronseparate* references (**Supplementary Figure S6b-c**).

**Figure 5.**
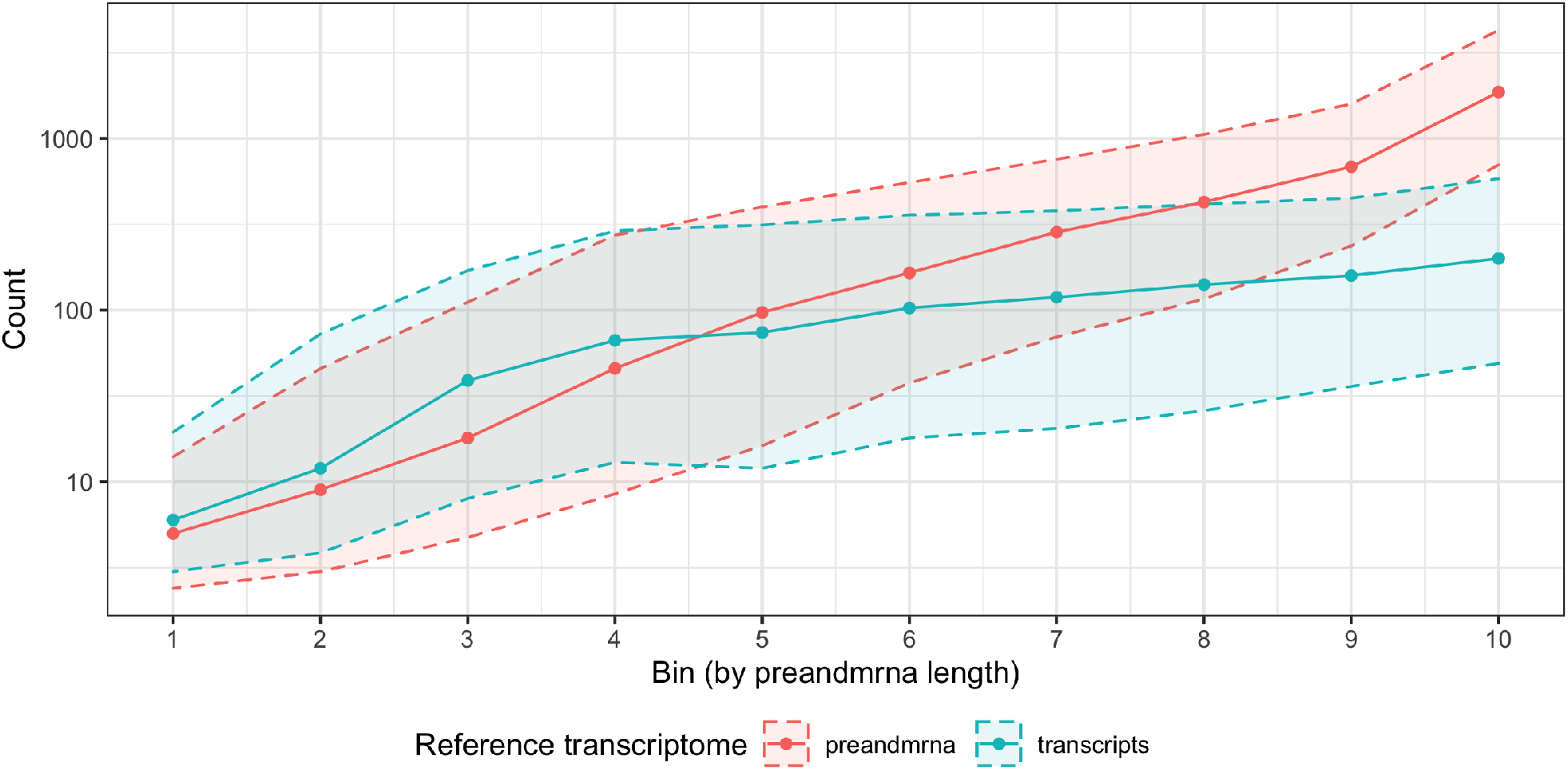
Chromium-based snRNA-seq data exhibit a gene length bias. Genes are binned by their length into ten equally-sized bins (*x*-axis) where the smallest bin number corresponds to the shortest genes and the largest bin number corresponds to the longest genes. Using gene counts derived from either the *preandmrna* (red) or *transcripts* (blue) reference transcriptome, the distribution of gene counts across nuclei (*y*-axis) are shown with the median (solid points), and the 25th and 75th percentile (dashed lines). Genes are binned (*x*-axis) using the full gene length with both exons and introns, referred to as the ‘preandmrna length’.

Next, we explore potential causes for this bias. One previously described mechanism that could explain the length bias is internal priming (Chamberlin, Quinlan, 2020; *Interpreting Intronic and Antisense Reads in 10x Genomics Single Cell Gene Expression Data* 2020; Svoboda et al., 2021). Here, the poly(dT) primer primes at an internal poly-A sequence rather than the poly-A mRNA tail. The end result is that a single transcript can erroneously get counted multiple times if there are multiple stretches of poly-A sequences. Now, internal poly-A sequences can theoretically occur in either exonic or intronic regions, but intronic regions are typically longer and thus more likely to contain internal poly-A sequences (Sakharkar et al., 2004). Thus, the effect of internal priming is likely to be stronger for intronic reads than exonic reads, which would then explain the result in Figure 5 with a greater length bias for the *preandmrna* reference than the *transcripts* reference.

However, we found that internal priming does not fully explain the observed length bias. Using the *preandmrna* reference (**Figure 6a**), we found that given the same preandmrna gene length, genes with at least one internal poly-A 8-mer have higher expression than genes without any poly-A 8-mers on average. This supports the idea that internal poly-A priming is driving at least some portion of the gene length bias. However, we also see that among genes that do not have any internal poly-A 8-mers, a length bias can still be observed (blue line). This result holds true for different poly-A *n*-mer cut-offs (poly-A 6-mers, 10-mers, and 12-mers) (**Supplementary Figure S7**).

**Figure 6.**
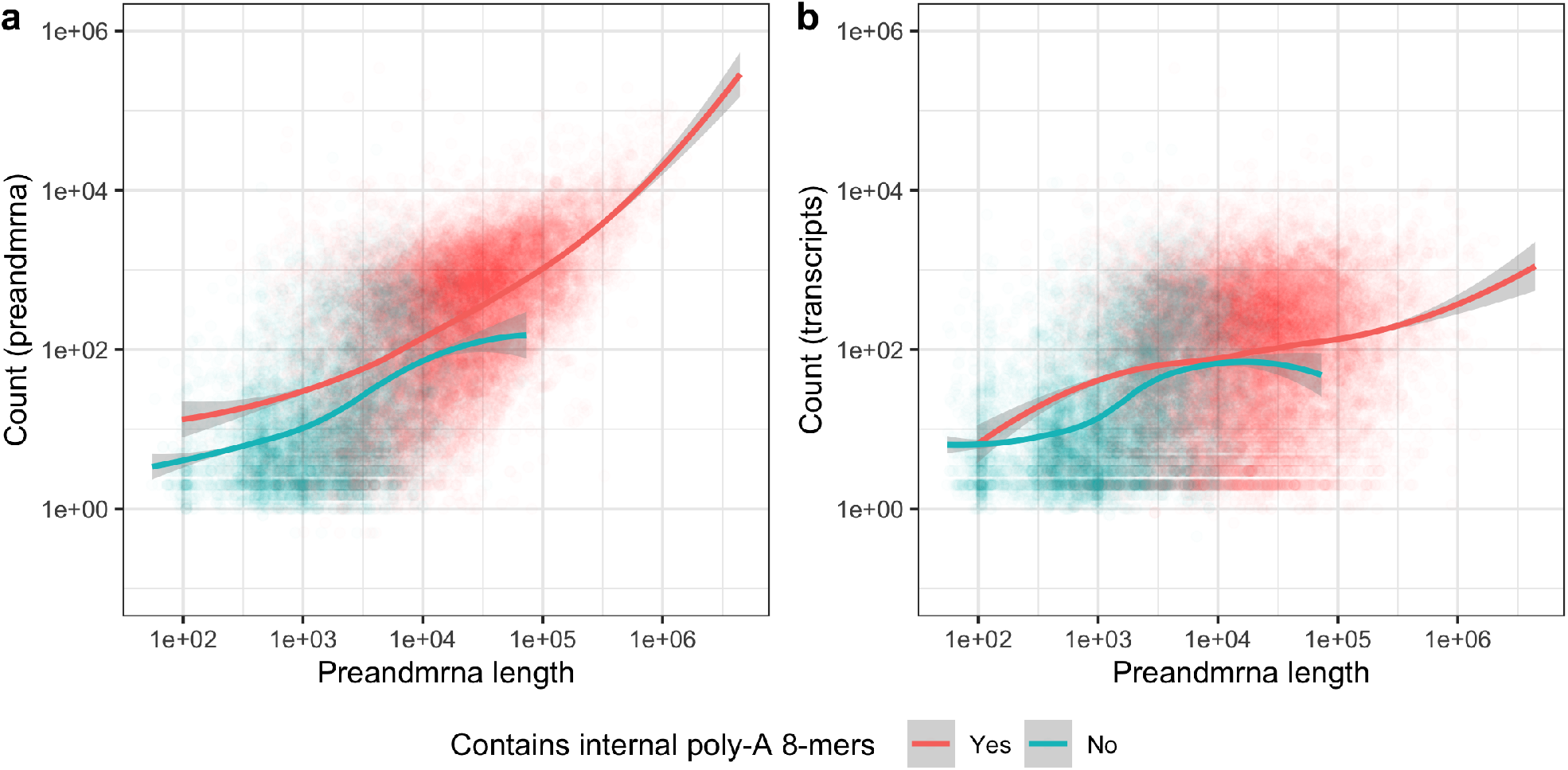
Comparison of gene length bias for genes with and without internal poly-A sequences. Each point is a different gene and are colored red if they have at least one internal poly-A 8-mer and blue if they do not and a loess curve is drawn for each set of genes. The *x*-axis uses the full gene length with both exons and introns (‘preandmrna’ gene length). The *y*-axis plots the sum of reads across all nuclei (base-10 log scale) from the **(a)** *preandmrna* reference or **(b)** *transcripts* reference.

Applying the same analysis with the *transcripts* reference (**Figure 6b**), we observe that first, in comparison to the *preandmrna* reference, there is less of a difference between genes with internal poly-A 8-mers and genes without internal poly-A 8-mers. This suggests that, as expected, internal priming explains less of the bias among the exonic reads. Second, for the genes without any internal poly-A 8-mers, there is still a clear length bias, similar to what we saw in the *preandmrna* reference. As the *transcripts* reference only includes exonic regions, this suggests that a significant portion of the length bias observed in the *preandmrna* reference that is not explained by internal priming actually resides in the exonic reads.

To further investigate mechanisms, we compared the strength of the bias when using the preandmrna length, where we include both intronic and exonic regions, versus the transcript length, where we only include the exonic regions. As the exonic region is a subset of the full gene, these two lengths are correlated (**Supplementary Figure S6a**), hence we would expect a length bias with both. The respective strengths of the bias, however, can tell us whether the causal mechanisms behind the bias is something we should expect to scale with the preandmrna length or the transcript length.

We found that the *preandmrna* reference exhibits a length bias that is more correlated with the preandmrna length than the transcript length. Under a base-10 log scale, the overall Pearson’s correlation coefficient between the counts and the preandmrna length is *r* = 0.68, while *r* = 0.37 between the counts and the transcript length (**Supplementary Figure S9a,b**). In comparison, with the *transcripts* reference, *r* = 0.39 between the counts and the preandmrna length and *r* = 0.38 between the counts and the transcript length (**Supplementary Figure S9c,d**). This suggests that part of the length bias lies with the intronic reads in the *preandmrna* reference and is correlated with the length of the intronic region, something that a mechanism like internal priming would be consistent with. However, there is another part of the length bias that lies with the exonic reads and is less correlated with either the preandmrna or transcript length. This reinforces our previous conclusion that there are likely multiple sources for the length bias, which may be different for intronic reads versus exonic reads.

## DISCUSSION

Droplet-based snRNA-seq technologies are becoming the preferred technology to profile gene expression in frozen cells or cells that are hard to dissociate. With these new data come new statistical challenges that need to be addressed, including how to model these data. Here, we demonstrate that Chromium-based snRNA-seq data are not zero-inflated and follow a negative binomial (NB) distribution. These data can also be approximated by binomial and Poisson distributions. Our results suggest that statistical methods that depend on these assumptions, such as tools for batch correction (Satija et al., 2015) or differential expression analysis (Robinson et al., 2010; Anders, Huber, 2010; Risso et al., 2018) commonly used for scRNA-seq, can likewise be used for snRNA-seq. As a general example, our results demonstrate that a NB generalized linear model *g*(*Y*) = *Xβ* can be used for snRNA-seq data, where *Y* are the counts and *X* are the variables of interest.

Furthermore, we show that choices in the reference transcriptomes used to perform quantification mapping of snRNA-seq data can impact both the fraction of reads mapped and downstream analyses, such as cell type classification. This is meaningful as different annotated cells can result in different biological interpretations of the same data. Standard quantification tools used for scRNA-seq are therefore not sufficient for analyzing snRNA-seq and the incorporation of intronic regions in the quantification of scRNA-seq data is an important consideration. In addition, we show that the choice of how intronic reads are included in quantification is less important than the choice to do so. Both in terms of library size and cell type classification, we found similar performance using reference transcriptomes that incorporate intronic regions.

With respect to cell type classification, we note that the higher agreement with cell type labels derived by the Ding et al., 2020 authors in the reference transcriptomes that incorporate intronic regions (*preandmrna, introncollapse*, and *intronseparate*) is not surprising given that they also include intronic regions in their quantification tool (Ding et al., 2020). However, we do not claim that concordance between the cell type labels imply that our assigned cell type labels are “correct.” Instead, we simply demonstrate that there are significant differences in cell type classification that arise from the choice to include or not include intronic regions in quantification. Since the disparities in mapping rate between references that do not include intronic regions (*transcripts*) compared to those that do (*preandmrna, introncollapse*, and *intronseparate*) lead to differences in counts that do not occur uniformly across all marker genes.

Across the references, we also observe a gene length bias in snRNA-seq. This bias is strongest when intronic regions are included, which may partly explain why a length bias has not been previously described with scRNA-seq data using the same sequencing technology. However, we also showed how the bias is not limited to intronic regions and is present in exonic regions, as well. Previously proposed mechanisms like internal priming do not fully explain the observed bias, particularly for exonic regions, and we leave further investigation of potential causes to future work.

Our work comes with limitations. First, as we use experimental mouse cortex nuclei data for our analysis, we can only create “pseudo-negative” control datasets that are not completely biologically homogeneous. For this reason, our results can be considered as a maximum bound on the amount of overdispersion. Since we have found that Chromium-based snRNA-seqdata follow a NB distribution and can be approximated by the binomial or Poisson distribution in many cases, we expect that the true measurement error should be even lower than what we have found. Negative controls of technical replicates have previously been generated to study technical variability in scRNA-seq data (Zheng et al., 2017) and a similar dataset for snRNA-seq can, in principle, be used to verify our conclusions.

Other limitations are that we only investigated one high-throughput experimental technology to capture gene expression (10x Genomics Chromium). We also only considered nuclei from two biological replicates (two mouse cortices) from one study. However, we do not expect our results on the distributions for measurement error of snRNA-seq dataset to change with droplet-based protocols with UMIs, as similar analyses for scRNA-seq found consistent results across three platforms (Svensson, 2020).

We note that the magnitude and nature of the effect of quantification choices on downstream analyses may vary depending on the dataset. As we observed in our dataset, some cell type classifications appear to be more affected by the inclusion of intronic reads than others. However, since we also observed large disparities in library size depending on whether intronic reads are included in the reference transcriptome and this phenomenon is likely to be agnostic to different snRNA-seq datasets, our results suggest that quantification choices will be informative for downstream analyses of snRNAseq data. Given the relative ease of including introns in quantification with tools like `salmon alevin` and the significant loss in information when they are not included, the inclusion of intronic reads is a crucial step in the analysis of snRNA-seq data.

## METHODS

### Data

The mouse whole cortex nuclei dataset was generated by Ding et al., 2020 using the 10x Genomics Chromium platform (Zheng et al., 2017). Two experiments were performed, resulting in two biological replicates (Cortex1 and Cortex2). Each of these experiments was run on a platform with two flow cells with four lanes each, resulting in eight SRA files for each experiment. After running `salmon alevin` (Srivastava et al., 2019) to map the reads with the different transcriptome indices described in the next section, we read in the nucleus × gene UMI count matrices as a `SingleCellExperiment` object using the `tximeta` (Love et al., 2020) and `SingleCellExperiment` R packages (A Lun, Risso, 2020).

### Quantification with four sets of reference indices

We started with Sequence Read Archive (SRA) data downloaded from the Gene Expression Omnibus with accession number GSE132044 and converted them into FASTQ files using the SRA toolkit. The FASTQ files, along with a reference transcriptome, are fed into `alevin` for quantification (Srivastava et al., 2019). To create the reference transcriptome, we processed Gencode reference files, in particular the GRCm38 primary assembly FASTA file and the Gencode vM25 gene annotation GTF file.

In total, four reference transcriptomes were created. Each of these reference transcriptomes differ in that they incorporate intronic regions into its target sequences in different ways. We define the following types of target sequences. First, we start with the transcript sequences, which are defined from the downloaded genome sequence and GTF file and consist of the exonic regions for each transcript. These are what we call “spliced transcripts” and each transcript is a separate target sequence. We can create “unspliced transcripts” as target sequences by re-adding the intronic regions between any two exonic regions in a transcript. The length of each unspliced transcript must therefore be greater than or equal to its corresponding spliced transcript. Lastly, intronic regions for a given transcript or gene can themselves be used as target sequences. Based on the work of Soneson et al., 2021, we define the introns in two ways: “separate” or “collapse.” In the “separate” approach, the intron target sequences are defined as the intronic regions from a transcript of a given gene. In the “collapse” approach, the intron target sequences are defined to be the intronic regions of a gene that are not exonic in any isoforms of the gene. Thus, while the “separate” approach allows for intron target sequences to overlap with exonic regions of other transcripts, the “collapse” approach does not. For all intronic target sequences, a flanking length of 50bp (read length) is also added to account for reads that map to exon/intron junctions.

The different target sequences used to create the reference transcriptome form the basis of our four distinct quantification reference transcriptomes, which are summarized in Table S1. In all four references, the complete genome sequence was also added to the reference transcriptome to create a decoy-aware transcriptome and minimize the spurious mapping of reads to intergenic regions.

### Preprocessing and quality control

For quality control, we use `perCellQCMetrics` from the `scater` R package (McCarthy et al., 2017). The quality control procedure removes nuclei with low library sizes or few expressed genes and discards genes with zero counts across all nuclei. After these steps, the number of genes number of nuclei is 27651 × 5612 for the *transcripts* reference, 31701 × 5680 for the *preandmrna* reference, 31670 × 5686 for the *introncollapse* reference, and 31661 × 5686 for the *intronseparate* reference. The quantification and preprocessing steps are summarized in **Figure 1**.

For exploratory analysis, we run principal component analysis (PCA) on the log normalized counts. The counts are normalized using size factors computed by `calculateSumFactors` from the `scran` R package (ATL Lun et al., 2016) and PCA was performed using the `scater` R package (McCarthy et al., 2017).

### Description of methods for distribution plots

We first remove the major sources of biological variation by subsetting the nuclei by cell type and biological replicate (**Figure S1**) in order to obtain a pseudonegative control dataset. Each nucleus × gene matrix subset is then downsampled to remove variability due to differences in sequencing depth and obtain comparable library sizes across nuclei.

Given a downsampled *m* × *n* matrix *M_t_* with *m* genes and *n* nuclei of a given cell type *t*, we calculate the following values. Let *x_ij_* be the number of reads for gene *i* and nuclei *j*. For every gene *i*, the empirical mean is defined as 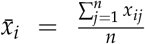, the empirical variance is defined as 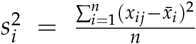, and the empirical probability or fraction of zero droplets is given by 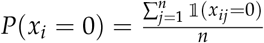.

The theoretical variances and probability of zero droplets is computed for each distribution using parameters estimated from the data. To estimate the parameters for a binomial distribution, *X_i_* ∼ *Binom*(*n, p_i_*), let *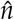* be the median column sum of *M_t_* and 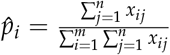. For a Poisson distribution, *X_i_*∼*Poisson*(*λ_i_*), let 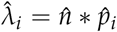. For a negative binomial (NB) distribution with an overall dispersion parameter, *X*_*i*_ ∼ *NB*(*ϕ, µ*_i_), where *ϕ* is the dispersion parameter and *µ_i_* is the mean, let 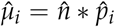. To estimate 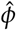, note that *σ*^2^ = *µ* + *µ*^2^/*ϕ*. Therefore, using the empirical means and variances for every gene *i*, we can estimate 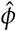 as the maximum likelihood coefficient from the following linear regression: 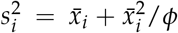. For a negative binomial distribution with gene-specific dispersion parameters, *X*_*i*_ ∼ *NB*(*ϕ_i_, µ_i_*), let 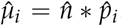. We estimate 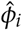 as the maximum likelihood estimate of a generalized linear model *g*(*E*(*X_i_*)) = *β*_0_ with the negative binomial family, where each observation is a different nucleus and a separate model is estimated for every gene using the `mgcv` package (SN Wood, 2017). This is what we refer to as the gene-specific (G-S) negative binomial distribution. After the parameters for each distribution have been estimated, we can compute the theoretical variances and probability of zero droplets, *P*(*X_i_* = 0), under each distribution. We also calculate the log-likelihood (LL) under each distribution for each gene.

The BIC log-likelihoods are computed using the formula BIC = *k* log(*n*) − 2 log(*L*), where *k* is the number of parameters, *n* is the number of observations, and *L* is the maximum likelihood (Schwarz, 1978). The BIC is calculated using the log-likelihood of the sum across all genes (assuming independence of genes) and observations. Code used to perform these computations is adapted from Townes et al., 2019 and all plots are generated using the `ggplot2` R package (Wickham, 2016).

For the goodness-of-fit tests, a Pearson’s chi-squared statistic was computed for every gene *i*. The formula for the Poisson distribution is given by 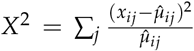 where the sum is over nuclei of a given cell type and biological replicate. 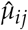 is the maximum likelihood estimate of the Poisson mean, and is given by 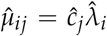 where 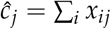 is the column sum for cell *j* and 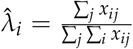 is the empirical rate at which reads maps to gene *i*. If the counts *x_ij_* are independent and follow a Poisson distribution with mean *µ_ij_*, then the statistics follow a chisquared distribution with *n* − 1 degrees of freedom (Marioni et al., 2008).

When *µ_ij_* is small (*µ_ij_ <* 1), as is often the case with snRNA-seq counts, the distribution of the chi-squared statistics is not well-approximated by the chi-squared distribution (G Wood, 2002). We found this in our application of snRNA-seq data as well (**Figure S8a**), where we ran the goodness-of-fit test on counts simulated from a Poisson distribution. We plot the quantile-quantile plots from a Poisson goodness-of-fit test for five different counts matrices, each with 21483 rows and 347 columns, which roughly corresponds to the number of genes (rows) and nuclei (columns) we encounter in our snRNA-seq dataset after restricting to a given cell type and cortex. Each counts matrix follows a Poisson distribution with a different mean parameter *µ*, ranging from 0.1 to 1.0. We observe that as *µ* decreases, the chi-squared statistics from the goodness-of-fit test increasingly deviate from the theoretical quantiles of a chi-squared distribution, which indicates that directly applying such a test to sparse counts matrices with low means is not a reliable test to assess the distributional fit.

To address this, we use grouped chi-squared tests following the method proposed by (G Wood, 2002). We first remove genes whose counts are too sparse and the number of cells we would need to group is more than what is available in our data. For the remaining genes, we use a grouped version of goodness-of-fit tests, where we first group the counts of *r* nuclei (G Wood, 2002). Let *y_ik_* = ∑*_j_ x_ij_* be the sum of the counts of the *r* nuclei in the *k*th group and let 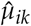 be the corresponding empirical mean for *y_ik_*. Since the sums of independent Poisson are also Poisson distributed, the chi-squared statistic follows a similar formula, 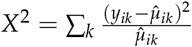, and is approximated by a chi-squared distribution with *n_k_* −1, where *n_k_* is the number of groups. We show that by applying this grouping procedure to simulated counts from a Poisson distribution, we get the expected results from the goodness-of-fit test (**Figure S8b**).

To determine the size of the group *r*, we choose 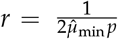, where 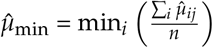, the smallest average empirical mean across genes, and *p* = 0.25. This ensures that the component variance of the chi-squared statistic is, on average, no larger than 2(1 + *p*), where 2 is the true theoretical variance of a 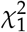 distribution.

For visualization purposes, the limits of the axes of the quantile-quantile plots are fixed to constant values across different cell types and reference transcriptomes.

### Cell type classification

For our analysis on distributions, we separate the mouse cortex nuclei using cell type labels computationally generated by (Ding et al., 2020). We compare these cell type labels to cell type labels generated using the `SingleR` R package and the built-in reference `MouseRNAseqData()` (Aran et al., 2019). Specifically, `SingleR` cell type labels are generated by comparing gene expression to the expression profile of the reference cells across marker genes. Each mouse cortex nuclei is assigned a cell type label based on similarity with Spearman correlation and labels are pruned by discarding ambiguous labels.

### Gene length bias

We define the gene length to be either the ‘preandmrna’ length, which is the full-length transcript with both exonic and intronic regions, or the ‘transcript’ length, which is the transcript with only exonic regions. For each gene, we calculate the sum of the expression counts across all nuclei in the dataset.

To count the number of poly-A sequences for each gene, we first count the number of poly-A n-mers in every transcript for a given gene. Then we summarize at the gene-level by taking the maximum number of poly-A n-mers across all transcripts (spliced and unspliced) for a given gene. This allows us to separate genes into two groups, that is, genes that do not have any poly-A n-mers for any of its transcripts and genes that have at least one poly-A n-mer in at least one of its transcripts.

## Code availability

The code used to produce our analyses is available at https://github.com/stephaniehicks/quantify-snrna.

## Supplementary Materials

### SUPPLEMENTAL TABLES

**Supplementary Table S1.** Summary of the reference transcriptome indices used in the quantification mapping tool. Reads are mapped to the target sequences in the reference transcriptome index. The reference transcriptome is augmented with decoy sequences, which mitigates the spurious mappings of reads that map better to the decoy sequences than the target sequences. In all reference transcriptomes, we provide the complete genome sequence as a decoy sequence to exclude reads coming from unannotated intergenic regions.

| Reference transcriptome | Target sequences | Decoy sequence |
| --- | --- | --- |
| *transcripts* | - spliced transcripts (mRNA): exonic regions of transcripts | genome |
| *preandmrna* | - spliced transcripts (mRNA): exonic regions of transcripts<br>- unspliced transcripts (pre-mRNA): full-length transcripts with both exonic and intronic regions | genome |
| *introncollapse* | - spliced transcripts (mRNA): exonic regions of transcripts<br>- introns “collapse”: intronic regions extracted after collapsing all transcripts of a gene | genome |
| *intronseparate* | - spliced transcripts (mRNA): exonic regions of transcripts<br>- introns “separate”: intronic regions extracted separately from each transcript isoform | genome |

**Supplementary Table S2.**
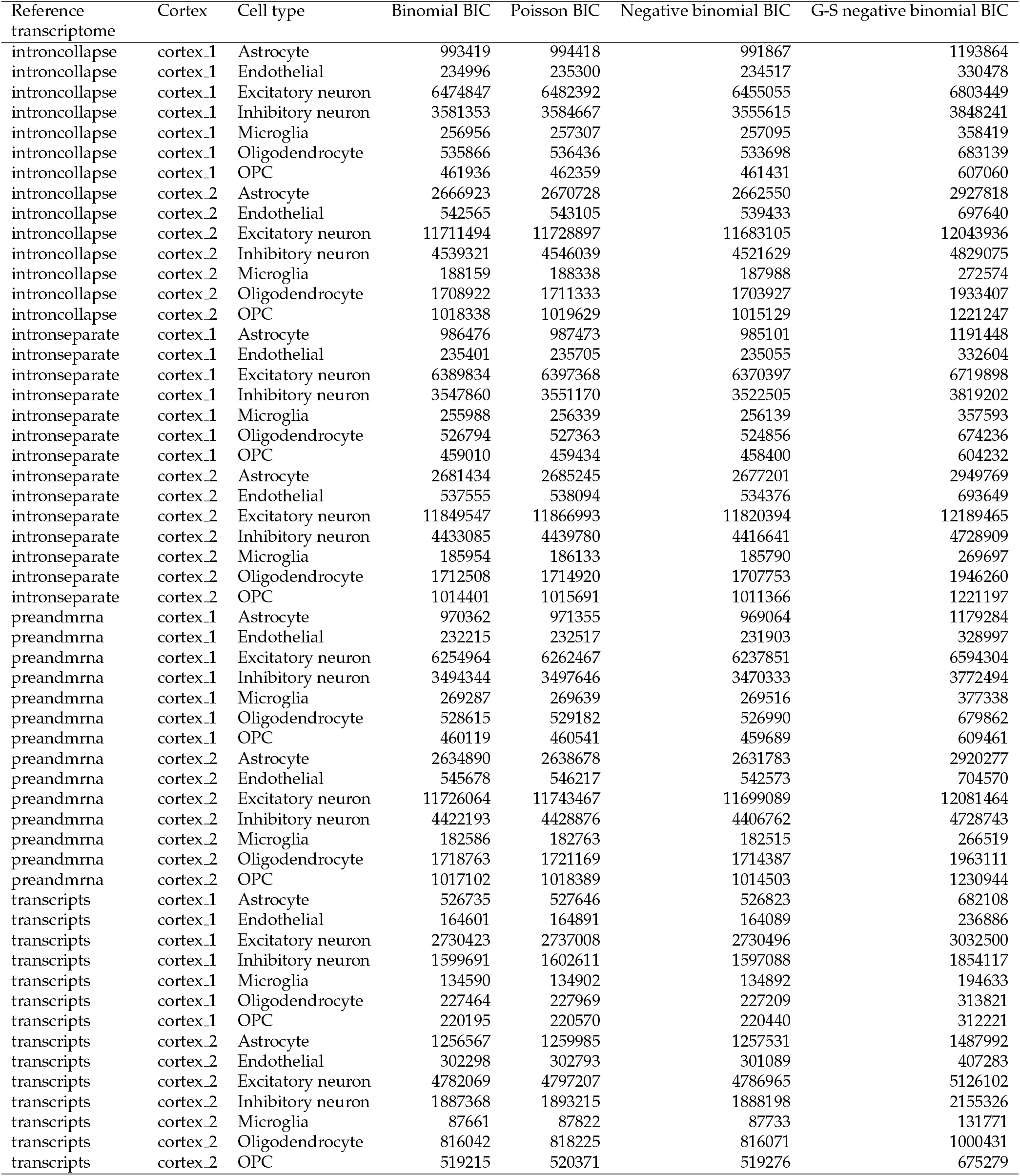
BIC log-likelihoods for each reference transcriptome, cortex, and cell type combination. The BIC log-likelihood are generally lowest for the negative binomial distribution or the binomial distribution, with similar BIC values for the Poisson distribution and higher values for G-S negative binomial, primarily due to the BIC penalty on the number of parameters. Negative binomial refers to a negative binomial distribution with one overdispersion parameter for all genes. G-S negative binomial refers to a negative binomial distribution with gene-specific overdispersion parameters.

### SUPPLEMENTAL FIGURES

**Supplementary Figure S1.**
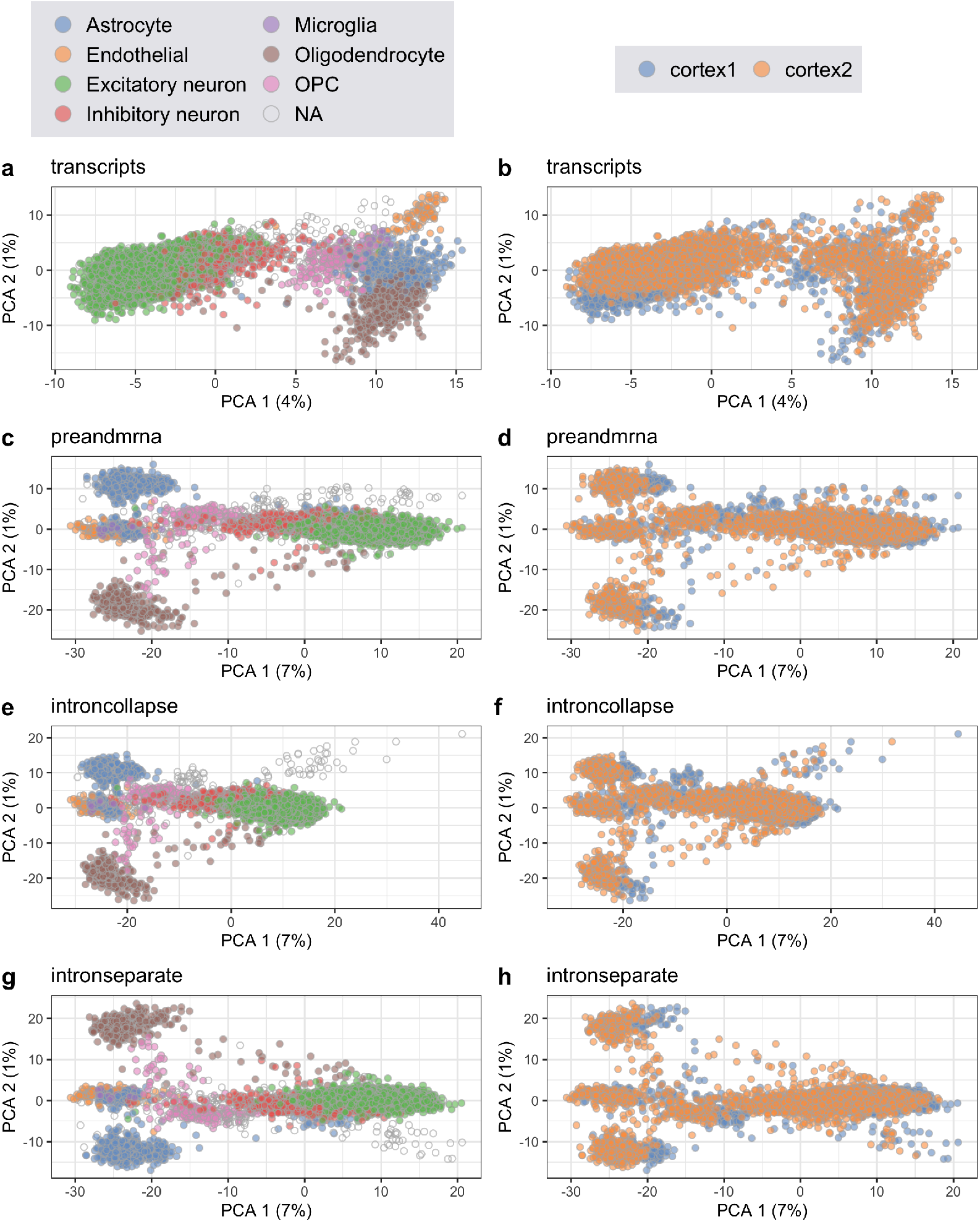
Principal components analysis to identify sources of biological variation among nuclei. The first two principal components from principal component analysis (PCA), which was performed on the normalized log-transformed counts using each reference transcriptome (rows). In the left column, we observe that in all references, the main source of variation is explained by the different cell types (colors are cell type labels as classified by Ding et al., 2020). In the right column, we also observe some minor variation by biological replicates (mouse cortices). **(a, b)** *transcripts* reference **(c, d)** *preandmrna* reference **(e, f)** *introncollapse* reference **(g, h)** *intronseparate* reference.

**Supplementary Figure S2.**
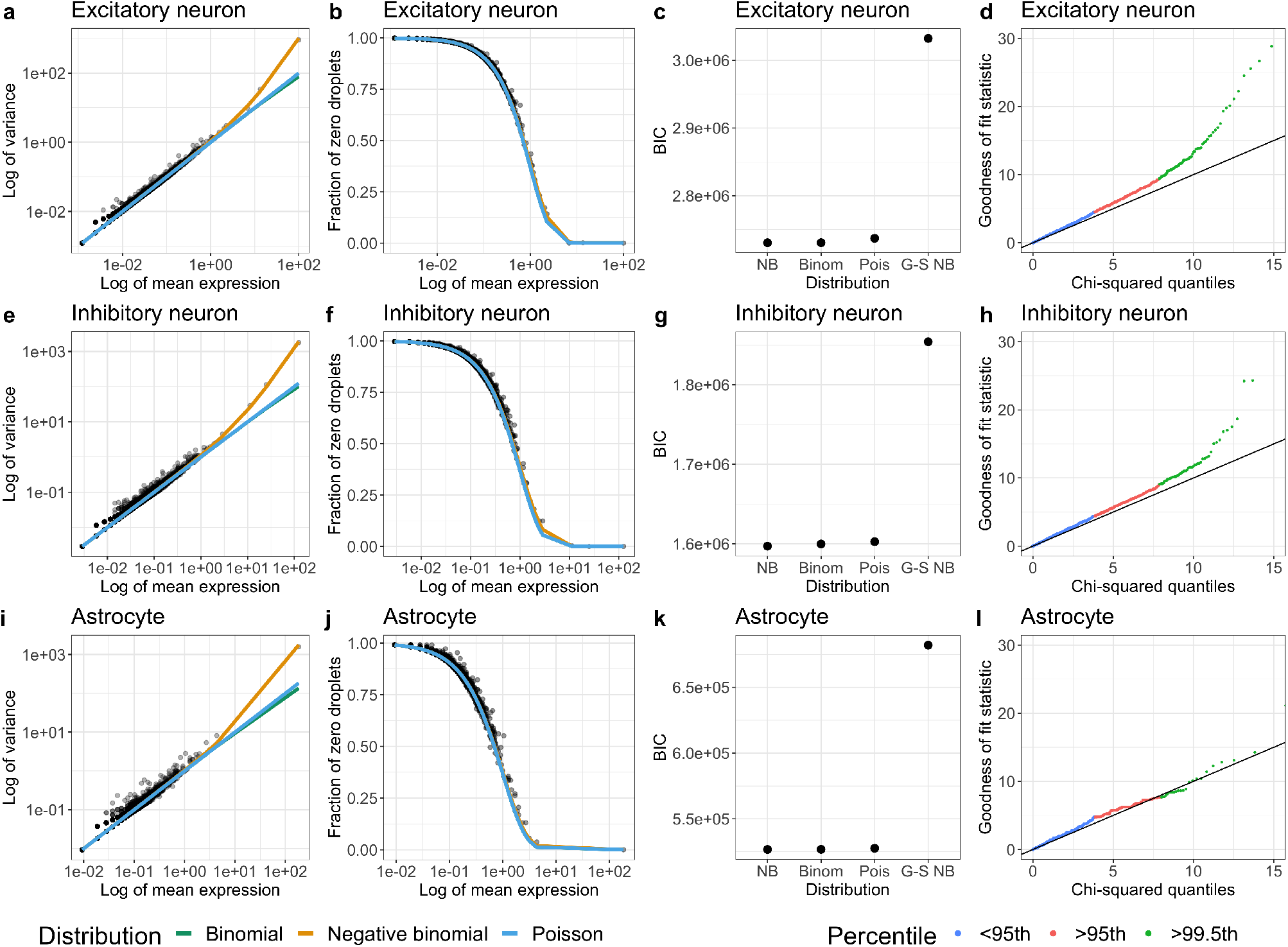
Chromium-based scRNA-seq data is not zero-inflated using the ‘transcripts’ reference. Similar to Figure 2 with subsets of cell types from Cortex 1, but using the *transcripts* reference transcriptome in the quantification mapping tool. **(a-d)** Excitatory neurons **(e-h)** Inhibitory neurons **(i-l)** Astrocytes.

**Supplementary Figure S3.**
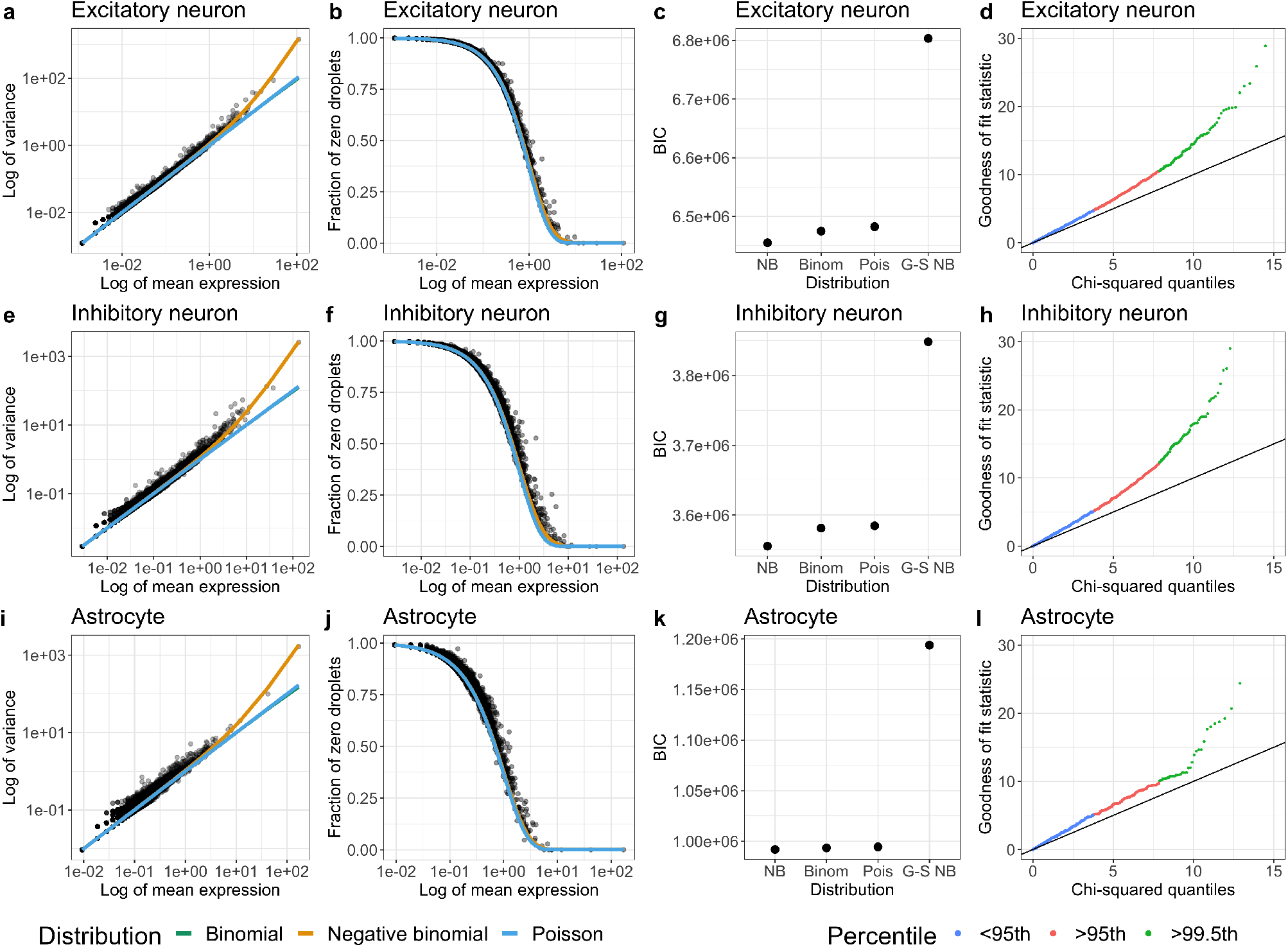
Chromium-based scRNA-seq data is not zero-inflated using the ‘introncollapse’ reference. Similar to Figure 2 with subsets of cell types from Cortex 1, but using the *introncollapse* reference transcriptome in the quantification mapping tool. **(a-d)** Excitatory neurons **(e-h)** Inhibitory neurons **(i-l)** Astrocytes.

**Supplementary Figure S4.**
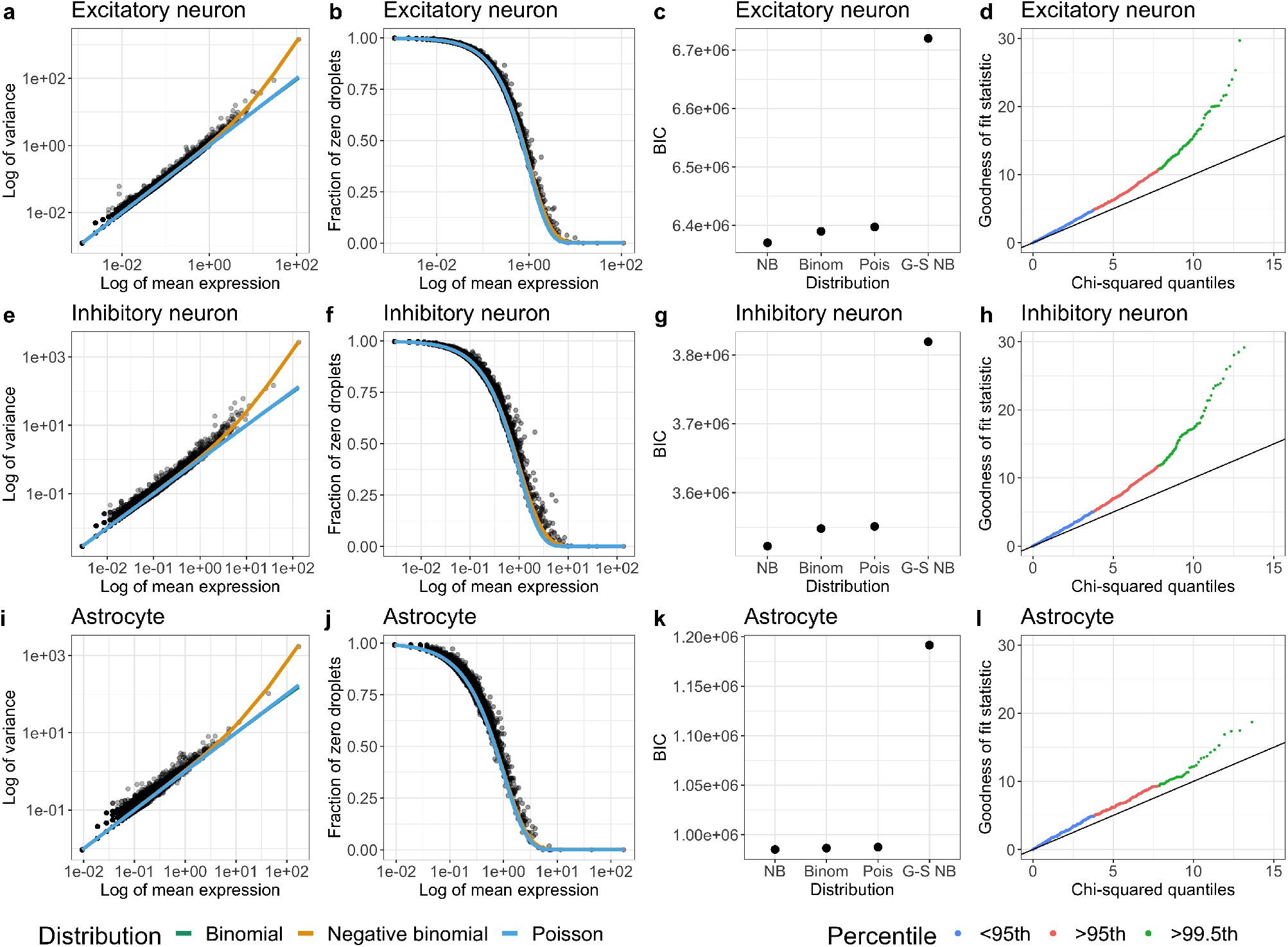
Chromium-based scRNA-seq data is not zero-inflated using the ‘intronseparate’ reference. Similar to Figure 2 with subsets of cell types from Cortex 1, but using the *intronseparate* reference transcriptome in the quantification mapping tool. **(a-d)** Excitatory neurons **(e-h)** Inhibitory neurons **(i-l)** Astrocytes.

**Supplementary Figure S5.**
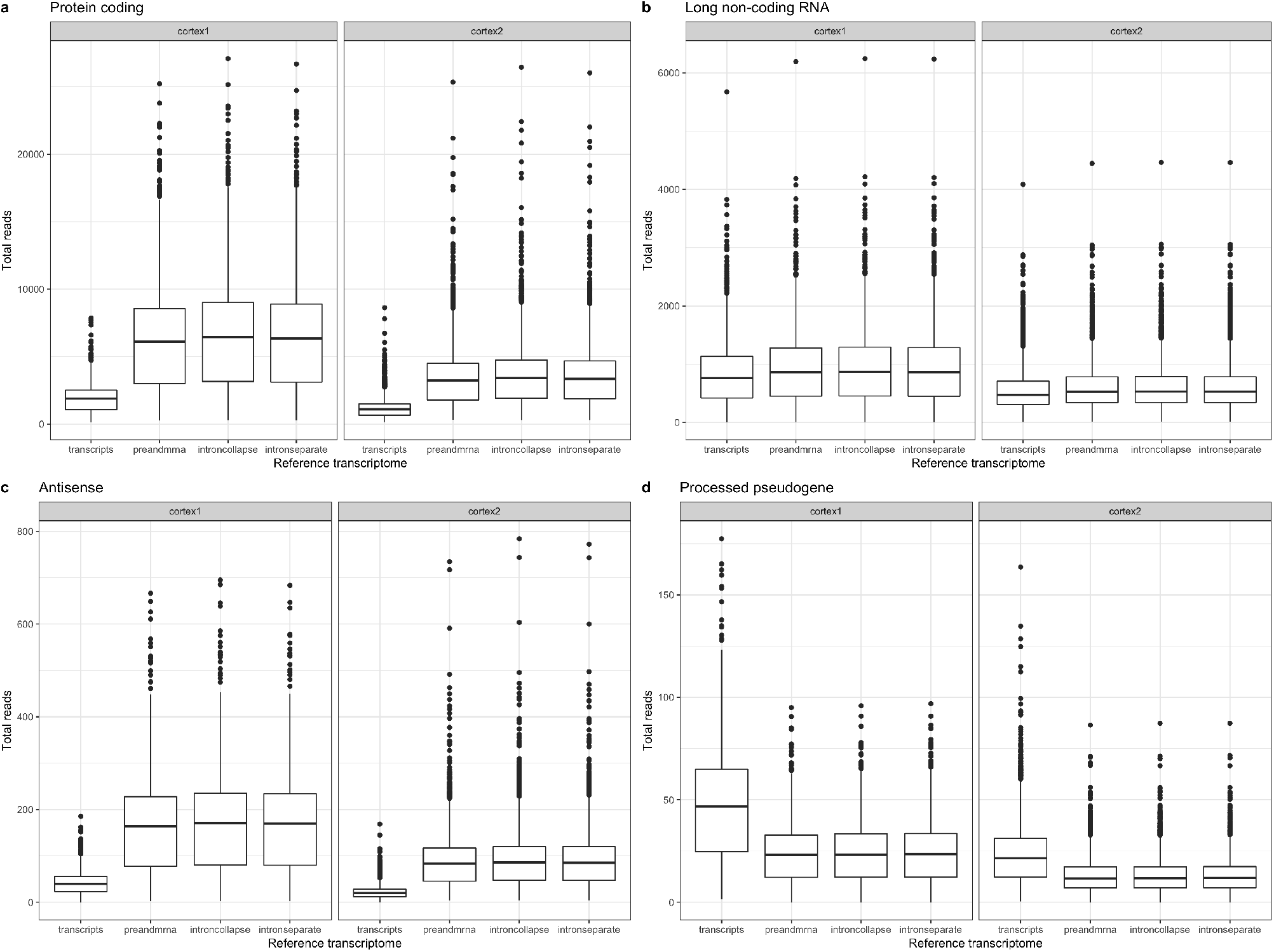
Number of mapped reads to gene sets stratified by reference transcriptomes. Similar to Figure 3, but now showing boxplots of the number of mapped reads to **(a)** protein coding genes, **(b)** long non-coding RNA, **(c)** antisense, and **(d)** processed pseudogene. For each gene type, the boxplots are faceted by the two biological replicates (Cortex1 and Cortex2).

**Supplementary Figure S6.**
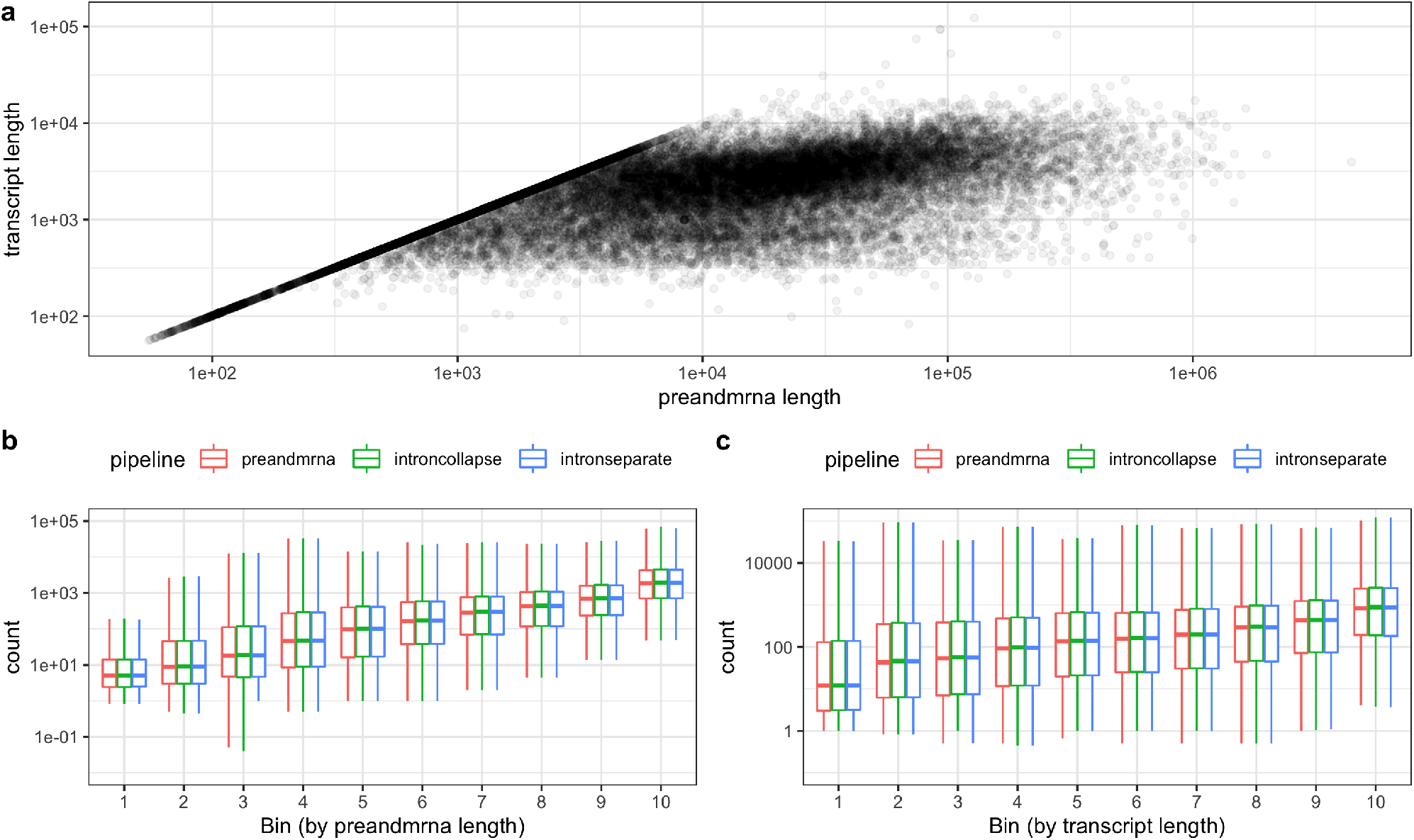
Correlation between preandmrna and transcript length and comparison of gene length bias across references with intronic reads. **(a)** For every gene, the preandmrna length is plotted on the *x*-axis and the transcript length is plotted on the *y*-axis. **(b-c)** A similar gene length bias is observed across the three references with intronic regions (outliers not included for boxplots). The sum of counts across all nuclei is plotted on the *y*-axis and genes are binned into ten equally-sized bins by their **(b)** preandmrna length or **(c)** transcript length.

**Supplementary Figure S7.**
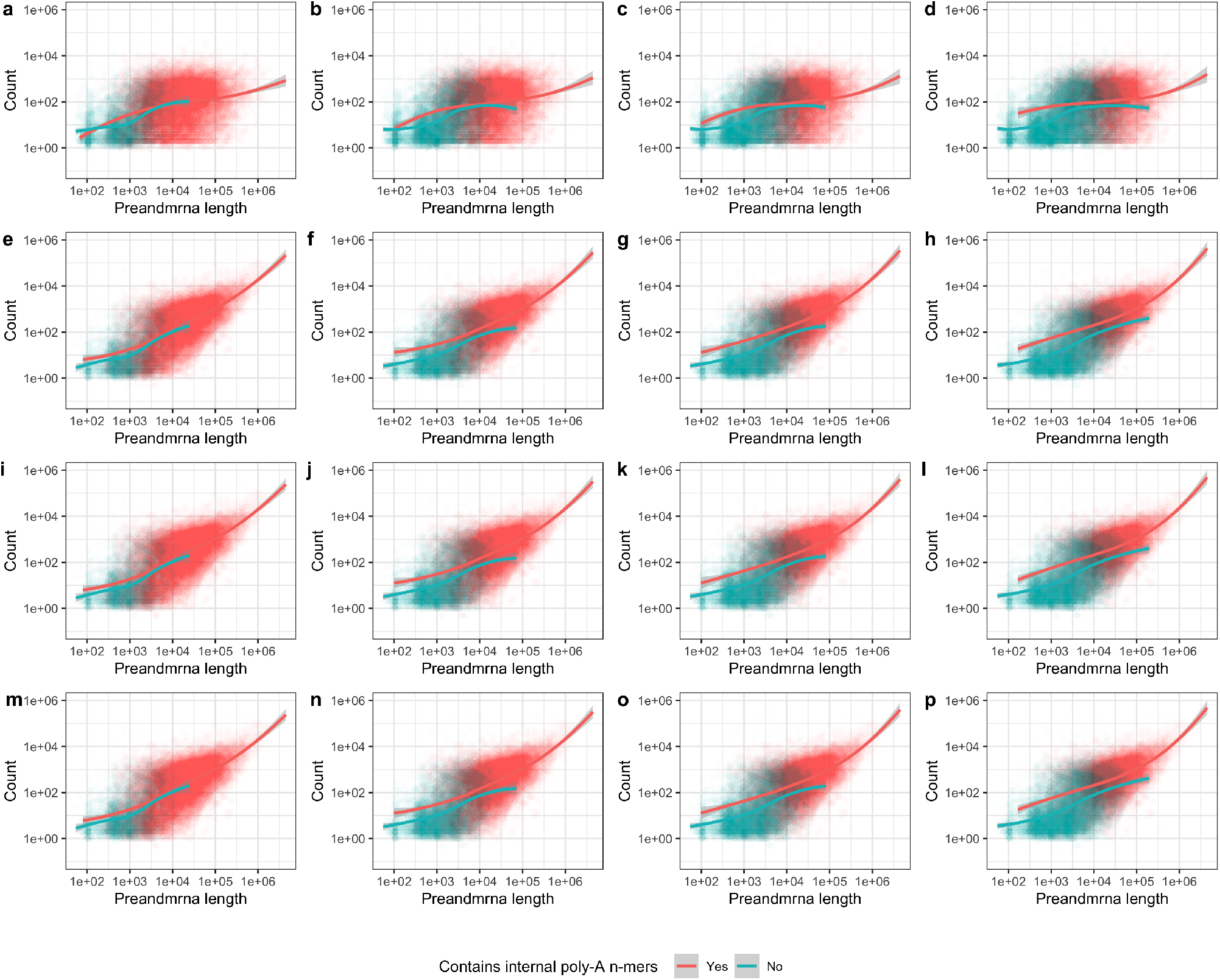
Comparison of gene length bias for genes with and without internal poly-A sequences under different reference transcriptomes and *n*-mer cut-offs. Each point is a different gene, with the sum of counts across all nuclei plotted on the *y*-axis (base-10 log scale). Genes are colored red if they have at least one internal poly-A *n*-mer and blue if they do not. A loess curve is drawn for each set of genes. The *x*-axis uses the full gene length with both exons and introns (‘preandmrna’ gene length). Each column corresponds to a different poly-A *n*-mer cut-off **(a, e, i, m)** poly-A 6-mer, **(b, f, j, n)** poly-A 8-mer, **(c, g, k, o)** poly-A 10-mer, **(d, h, l, p)** poly-A 12-mer. Each row corresponds to a different reference transcriptome **(a, b, c, d)** *transcripts*, **(e, f, g, h)** *preandmrna*, **(i, j, k, l)** *introncollapse*, **(m, n, o, p)** *intronseparate*.

**Supplementary Figure S8.**
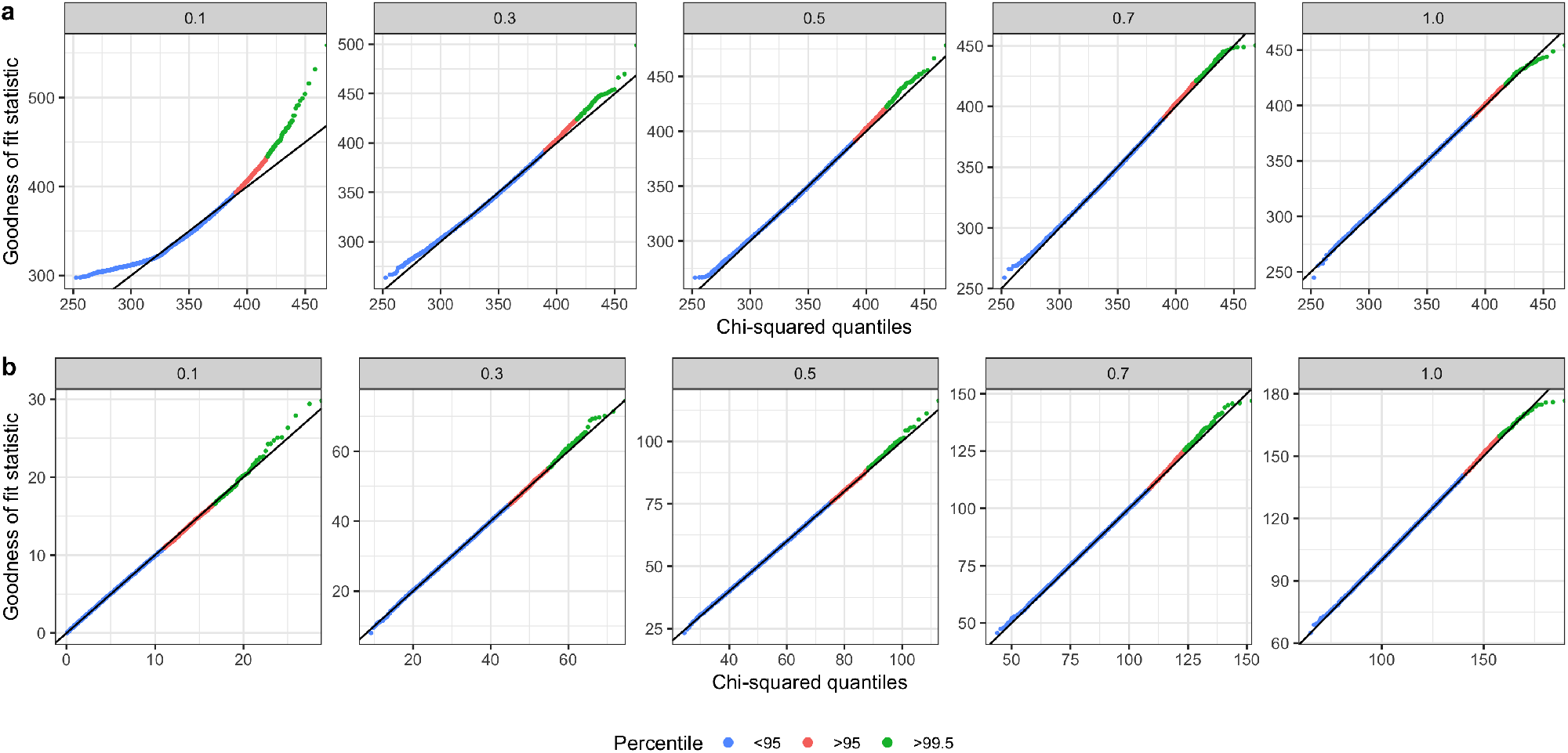
Comparison of grouped versus ungrouped Poisson chi-squared tests. Using matrices of simulated Poisson counts with different *µ* parameters (*µ* = 0.1, 0.3, 0.5, 0.7, 1.0 from left to right), the quantile-quantile plots from an **(a)** ungrouped Poisson chi-squared test and **(b)** grouped Poisson chi-squared test are compared.

**Supplementary Figure S9.**
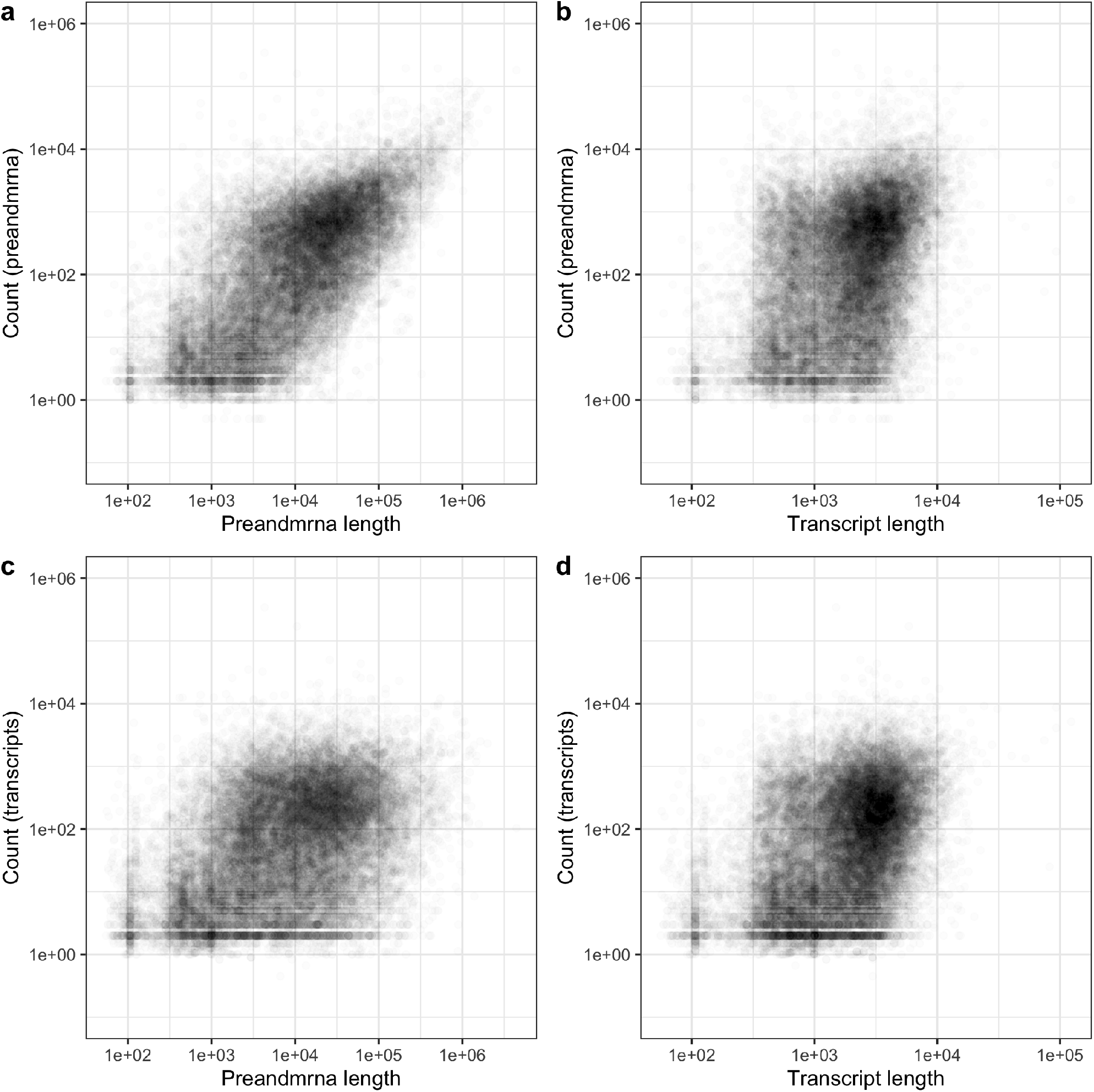
Comparison of correlations between length and counts for the *preandmrna* and *transcripts* references. Each point is a different gene, with the length (base-10 log scale) on the *x*-axis and the counts (base-10 log scale) on the *y*-axis. The length is defined as either the full gene length with both exons and introns (‘preandmrna’ length) or the length with only exons (‘transcripts’ length). The counts are defined as the sum of reads across all nuclei under a given reference transcriptome (*preandmrna* or *transcripts*). Pearson’s correlation coefficient (*r*) is calculated for each scatter plot. **(a)** ‘preandmrna’ length and *preandmrna* reference (*r* = 0.68) **(b)** ‘transcript’ length (*r* = 0.37) and *preandmrna* reference **(c)** ‘preandmrna’ length and *transcripts* reference (*r* = 0.39) **(d)** ‘transcript’ length and *transcripts* reference (*r* = 0.38).

**Supplementary Figure S10.**
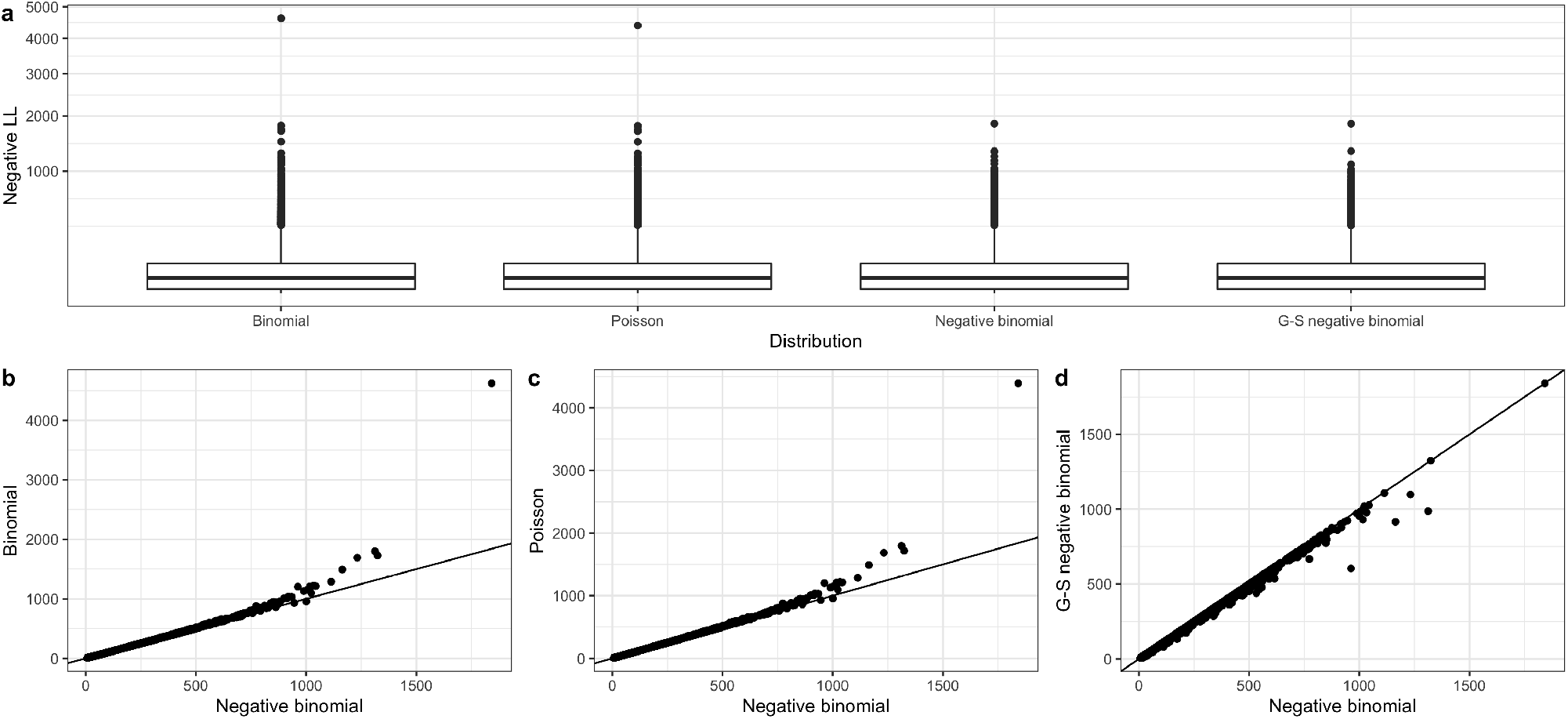
Assessment of probability distribution fits for each gene using negative log-likelihoods. For each gene (a dot in either the box plot or the scatter plot), the negative log-likelihood (LL) under each distribution is calculated and compared. **(a)** Box plots of the negative log-likelihoods for each distribution **(b)** The negative log-likelihood for every gene under the negative binomial distribution (*x*-axis) versus the binomial distribution (*y*-axis) **(c)** The negative log-likelihood for every gene under the negative binomial distribution (*x*-axis) versus the Poisson distribution (*y*-axis) **(d)** The negative log-likelihood for every gene under the negative binomial distribution (*x*-axis) versus the negative binomial distribution with gene-specific overdispersion parameters (*y*-axis). For each gene (black dot), if the negative LL fit is the same for each distribution, we expect it to fall along the black line (*y*=*x*).

